# Granulomatous inflammatory responses are elicited in the liver of PD-1 knockout mice by *de novo* genome mutagenesis

**DOI:** 10.1101/2023.08.17.553694

**Authors:** Ilamangai Nagaretnam, Azusa Yoneshige, Fuka Takeuchi, Ai Ozaki, Masaru Tamura, Shiori Suzuki, Toshiaki Shigeoka, Akihiko Ito, Yasumasa Ishida

## Abstract

**Aims:** programmed death-1 (PD-1) is a negative regulator of immune responses. Upon deletion of PD-1 in mice, symptoms of autoimmunity developed only after they got old. In a model experiment in cancer immunotherapy, PD-1 was shown to prevent cytotoxic T lymphocytes from attacking cancer cells that expressed neoantigens derived from genome mutations. Furthermore, the larger number of genome mutations in cancer cells led to the more robust anti-tumor immune responses after the PD-1 blockade. In order to understand the common molecular mechanisms underlying these findings, we hypothesize that we might have acquired PD-1 during evolution in order to avoid/suppress autoimmune reactions against neoantigens derived from mutations in the genome of aged individuals. Main methods: to test the hypothesis, we introduced random mutations into the genome of young PD-1^-/-^ and PD-1^+/+^ mice. We employed two different procedures of random mutagenesis: administration of a potent chemical mutagen N-ethyl-N-nitrosourea (ENU) into the peritoneal cavity of mice and deletion of *MSH2*, which is essential for the mismatch-repair activity in the nucleus and, therefore, for the suppression of accumulation of random mutations in the genome.

**Key findings:** we observed granulomatous inflammatory changes in the liver of the ENU-treated PD-1 knockout (KO) mice, but not in the wild-type (WT) counterparts. Such lesions also developed in the PD-1/MSH2 double KO mice, but not in the MSH2 single KO mice. Significance: the results we obtained support our hypothesis: PD-1 probably functions to avoid/suppress inflammatory responses against neoantigens derived from genome mutations in aged individuals.

## Introduction

Programmed death-1 (PD-1) was discovered in cells undergoing apoptosis [1]. PD-1 is expressed on antigen-activated T lymphocytes (T cells) and interacts with one of its ligands, programmed death-ligand 1 (PD-L1). The interaction causes phosphorylation of immunoreceptor tyrosine-based switch motif (ITSM), located in the cytoplasmic portion of PD-1, that further recruits SHP-2 tyrosine phosphatase to mediate the inhibitory function of the T cells [2]. To date, the PD-1: PD-L1 pathway has been shown to be exploited by our physiological self-components, microorganisms, and tumor cells to attenuate the T-cell responses [3].

The global absence of PD-1 in mice led to different types of immunological disorders based on the mouse’s genetic background [4,5]. Autoimmune diseases that develop spontaneously in PD-1^-/-^ mice, particularly those on the C57BL/6 (B6) genetic background, are unique because the phenotype observed are very mild and late-onset [4].

In a model experiment of cancer immunotherapy, PD-1 was shown to prevent cytotoxic T cells that could recognize and respond to mutant antigens (in the context of a class I MHC molecule) from attacking cancer cells with the cognate genome mutations [6]. Interestingly, the larger number of genome mutations in cancer cells led to the more robust immune responses against such cells, but only after the PD-1 blockade [7].

In order to understand the common molecular mechanisms involved in these findings, a hypothesis about the physiological function(s) of PD-1 was presented [8]: When we are young, somatic cells in our bodies are pretty intact and harbor only a limited number of genome mutations. As we get old, however, *de novo* mutations are gradually accumulated in the genome of our somatic cells, and our highly sophisticated immune system with remarkable sensitivity and specificity could detect and misrecognize the resultant neoantigens as ‘nonself’, mounting the attacks against such aged somatic cells. We might have acquired PD-1 during evolution in order to avoid/suppress autoimmune reactions against gradually accumulating neoantigens in our aged somatic cells. When PD-1 suppresses harmful immune responses against normal somatic cells harboring neoantigens encoded by genome mutations in aged individuals, the beneficial immunity against neoantigens generated in cancer cells also should be suppressed by PD-1 [8].

To test the above hypothesis about the physiological function(s) of PD-1, we tried to introduce random mutations into the genome of somatic cells in relatively young PD-1^-/-^ and PD-1^+/+^ mice. We employed two different procedures of random mutagenesis of somatic cells: administration of a potent chemical mutagen N-ethyl-N-nitrosourea (ENU) into mice [9,10,11] and deletion of *MSH2*, which is essential for the mismatch-repair activity in the nucleus and, therefore, for the suppression of accumulation of random mutations in the mouse genome [12,13,14,15].

If the above hypothesis about the physiological function(s) of PD-1 is true, we should be able to observe some abnormal immune reactions against the mutation-derived neoantigens in PD-1^-/-^ mice but not in PD-1^+/+^ mice. We believe our current data shown here strongly support the idea [8].

## Materials and Methods

### Mice

All mice analyzed in this study had the genetic background of C57BL/6N (B6N). The wild-type B6N mice were purchased from CLEA Japan. All the mouse colonies were maintained in a specific pathogen free breeding facility in NAIST. All of the animal maintenance and animal experiments performed for this study were approved by the Committee on Animal Research at NAIST. All methods were performed based on the Policy on the Care and Use of Laboratory Animals at NAIST.

### Administration of the ENU mutagen to PD-1^+/+^ and PD-1^-/-^ mice

PD-1^-/-^ mice [4] were obtained from Tasuku Honjo at Kyoto University (Kyoto, Japan). In Mouse Phenotype Analysis Division of RIKEN Bioresource Research Center (Tsukuba, Japan), PD-1^+/+^ and PD-1^-/-^ mice were injected twice intraperitoneally with a dose of 85 mg/kg of ENU at 8 and 9 weeks of age. After injection, these mice were transported and kept in the animal facility in NAIST for further observation. At three, four, five, and seven months of age, the ENU-untreated and ENU-treated mice (PD-1^+/+^ and PD-1^-/-^) were dissected. Upon dissection, the mouse organs were fixed in 4 % paraformaldehyde (PFA) in phosphate-buffered saline (PBS) overnight at 4 °C. The next day, all the collected mouse organs were processed for histopathological analysis.

### Breeding scheme of PD-1^-/-^ MSH2^-/-^ mice

The MSH2^+/-^ mice [14] were obtained from Teruhisa Tsuzuki at Kyushu University (Fukuoka, Japan). For the control study, PD-1^+/+^MSH2^+/-^ mice were crossed with PD-1^+/+^MSH2^+/-^ mice to produce PD-1^+/+^MSH2^+/+^ and PD-1^+/+^MSH2^-/-^ mice. For the production of double KO mice, the PD-1^-/-^ mouse was crossed with the MSH2 hetero (MSH2^+/-^) mice. The resulting progenies (PD-1^-/-^MSH2^+/-^ mice) were further crossed to produce PD-1^-/-^MSH2^-/-^ and PD-1^-/-^MSH2^+/+^ mice. At seven months after birth, PD-1^+/+^MSH2^+/+^, PD-1^+/+^MSH2^-/-^, PD-1^-/-^MSH2^+/+^, and PD-1^-/-^ MSH2^-/-^ mice were dissected. Upon dissection, the mouse organs were fixed in 4 % PFA/PBS overnight at 4 °C. The next day, all the collected mouse organs were processed for histopathological analysis.

### Genotyping PCR of PD-1^-/-^ MSH2^-/-^ mice

Genomic DNA was extracted by using an alkaline method from the ear or tail of mice. The ear or tail pieces were lysed in 150 μl of the alkaline solution containing 10 N NaOH and 0.5 M EDTA at 100 °C for 30 min. Then, 150 μl of the neutralizing solution, 1 M Tris-HCl (pH 7.5), was added and mixed. The genomic DNA was centrifuged and kept at -20 °C for further analysis. PCR primers used for genotyping are shown in the table below. The thermal cycling condition with the KOD-FX enzyme (TOYOBO) for the detection of the PD-1 alleles was as follows: initial denaturation at 94 °C for 2 min; followed by 33 cycles of 94 °C for 30 sec, 64 °C for 30 sec, and 72 °C for 60 sec; then a final extension at 72 °C for 300 sec. This produces the 207 base-pair (bp) and 125 bp bands for the KO and wild-type WT alleles, respectively. The thermal cycling condition with KOD-FX for the detection of the MSH2 alleles was as follows: 94 °C for 2 min; followed by 35 cycles of 98 °C for 10 sec, 68 °C for 90 sec. This produces the 725 bp and 464 bp bands for the WT and KO alleles, respectively.

## Multiplex PCR primers used for the genotyping of KO mice

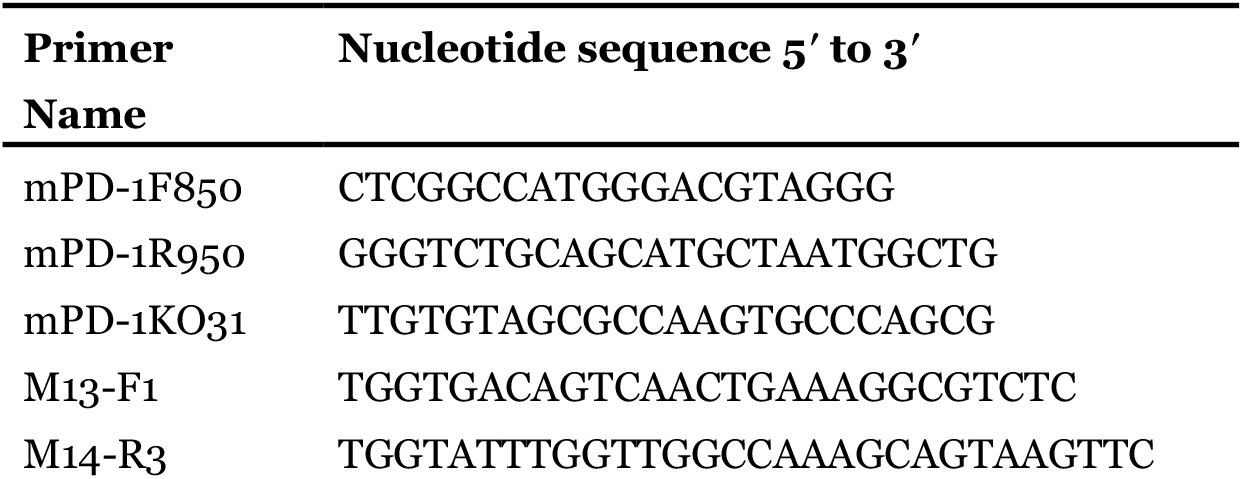
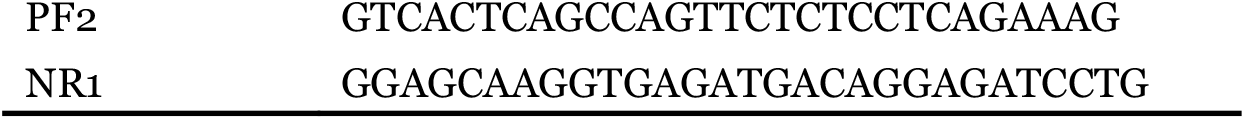

### Tissue preparation for histologic and immunofluorescence examination

Upon dissection and fixation with 4 % PFA, tissues were dehydrated through 50 % ethanol in PBS for 1 hour, followed by 70 % ethanol in DDW and kept for long-term storage prior to tissue processing. Tissues were further dehydrated through a graded series of ethanol by a tissue processor (Tissue-Tek VIP). The mouse tissues were embedded into paraffin (Tissue prep, Fisher Scientific, T580) by a tissue embedder (Tissue-Tek TEC) and directly chilled on a paraffin cooling module (Tissue-Tek TEC). Three μm sections were then made from the pre-chilled paraffin block using a microtome (Microm HM 340 E).

After harvesting the mouse organs, fresh tissue samples were cut and embedded directly in the OCT compound (Sakura Tissue-Tek O.C.T.) to prepare frozen blocks. The tissue sample was then snap frozen into liquid nitrogen and stored at -80 °C for long-term use. Five μm sections were prepared from the frozen block at -15 °C using a cryostat (CryoStar NX70).

### Hematoxylin and eosin staining

The tissue sections were stained with hematoxylin and eosin solution for histological assessments. The paraffin sections were first dewaxed in xylene for 15 min and rehydrated with a series of ordered ethanol (100 %, 90 %, 80 %, and 70 %) for 3 min each. Then, the sections were washed once with distilled water for 5 min. Next, these samples were further stained in Mayer’s Hematoxylin (Wako) for 15 min and washed under running tap water for 7 min. Following this, the sections were stained with eosin Y for 45 sec and dehydrated using 80 %, 90 %, and 100 % graded ethanol series. Next, xylene was used for 15 min to clear the tissue sections. Then, the tissue sections were mounted with PathoMount (Fujifilm Wako). Finally, the mounted tissue sections were imaged and analysed using Nikon Digital Sight Ds-fi1 Microscope Cmount Camera. The paraffin blocks and slides mounted with stained tissues were kept at room temperature for long-term usage.

### Immunofluorescence staining

The tissue sections were subjected to acetone fixation, followed by blocking with 5 % Normal Goat Serum (Abcam) in D-PBS (-) to block the nonspecific binding sites for 1 hour. The primary antibody staining was diluted in D-PBS (-) and stained overnight at 4 °C. The tissue samples were then washed with 1X PBS followed by fluorochrome-conjugated secondary antibody staining diluted in D-PBS (-) for one hour at room temperature. The tissue samples were then washed with 1X PBS. Next, the nuclei were stained with 4’,6-diamidino-2-phenylindole (DAPI), and the tissue sections were mounted with Immunoselect Antifading Mounting Medium (DNV Dianova).

Immunofluorescence staining was performed using primary antibodies: Armenian hamster anti-mouse CD3 (clone 145-2C11, eBioscience) and rat anti-mouse CD11b (clone M1/70, eBioscience). The CD3^+^ primary signal was amplified with an anti-Armenian hamster secondary antibody conjugated to Alexa Fluor 647, and CD11b^+^ primary signal was amplified with an anti-rat secondary antibody conjugated to Alexa Fluor 555. Finally, samples were imaged using confocal laser scanning microscope LSM 710 (Carl Zeiss) at 200x magnification and further processed using ImageJ (National Institutes of Health).

### Statistical analysis

The hepatocytic necrosis, lymphocytes infiltration, granulomas, and total inflammatory foci were quantitatively measured using the NIS Elements software (NIKON). *N* indicates the number of mice per group. Data are expressed as mean ± standard error of the mean (SEM). Statistical significance was calculated in RStudio (R-4.1.0) using the two-tailed Mann-Whitney *U* Test method. Statistical significance was defined as *P < 0.05.

## Results

### ENU promotes formation of the granulomatous lesions in the liver of PD-1^-/-^ mice

In mice, a major mutation spectrum caused by ENU has been observed between adenine-thymine pairs (in which more than 80 % of the nucleotide changes have been identified) [9]. Such mutations usually distribute to both coding and noncoding sequences [10]. ENU does not require metabolic activation to generate and develop potential disease phenotypes [11]. Currently, ENU is believed to be one of the most efficient chemical mutagens that can cause mutations in the mouse genome [9].

We established a mutation-acceleration/accumulation model by introducing ENU into the genome of young PD-1^-/-^ mice. Interestingly, the ENU-treated PD-1^-/-^ mice induced the formation of granulomatous lesions (Figure 1), that are frequently observed in the chronic inflammatory human diseases with unknown etiology (*e.g*., sarcoidosis and Crohn disease) or in the chronic infection of humans with persistent microbes (*e.g*., tuberculosis) [26]. The granulomatous inflammation in this study was predominantly found at four months after birth (two months after ENU treatment) in the liver of PD-1^-/-^ mice. The lesions were observed either around the vein (Figure 1G) or in the liver parenchyma regions (Figure 1H) of ENU-treated PD-1^-/-^ mice. These results suggest that the accumulation of genome mutations accelerated by ENU results in the development of granulomas in young PD-1^-/-^ mice.

**Figure 1.**
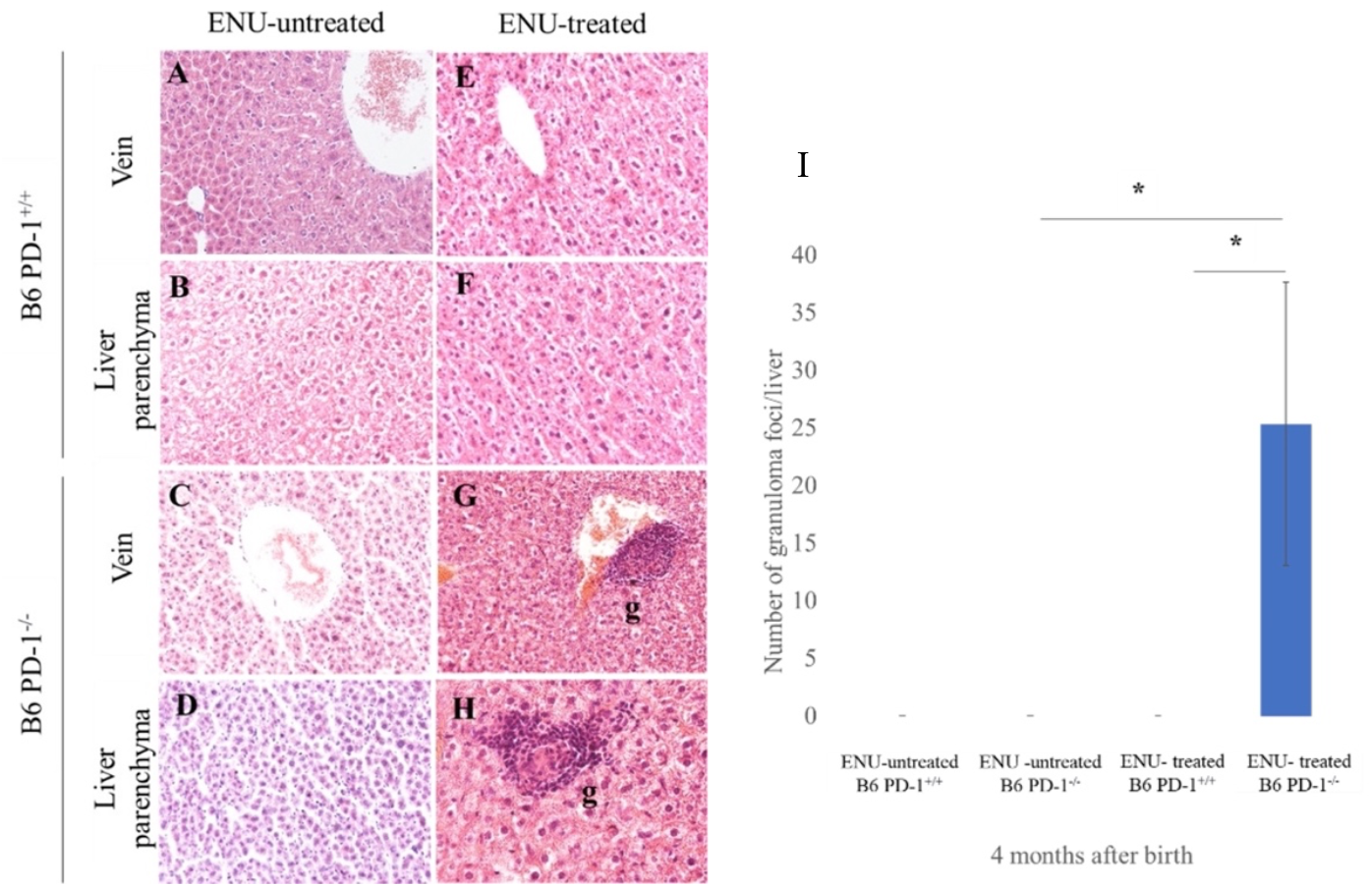
ENU promotes formation of the granulomatous lesions in the liver of PD-1^-/-^ mice. (**A-H**) Representative histological sections of liver tissues from ENU-untreated PD-1^+/+^ (**A-B**) and ENU-untreated PD-1^-/-^ mice (**C-D**), ENU-treated PD-1^+/+^ (**E-F**) and PD-1^-/-^ mice (**G-H**) at four months of age (two months after ENU treatment) are shown. Liver sections were examined by hematoxylin and eosin staining (200x magnification). Granulomas were seen in the liver tissue in both vein (**G**) and liver parenchyma (**H**) regions of ENU-treated B6 PD-1^-/-^ mice. Aggregation of lymphocytes surrounding macrophages exhibits the formation of granulomatous lesions (**G-H**). g(s) represent granulomas (**G-H**). (**I**) The bar graph showed number of granuloma foci from ENU-untreated and ENU-treated mice (PD-1^+/+^ and PD-1^-/-^). Four (right lateral, left lateral, right medial, and left medial) liver lobes in every mouse were used in the analysis (*N* ≥ 2 mice per ENU-untreated and *N* ≥ 4 mice per ENU-treated group). Data are expressed as means ± SEM. Statistical significance was determined by a two-tailed Mann-Whitney *U* test. *P<0.05.

The immunofluorescence analysis was carried out to detect the types of immune cells inside the inflammatory lesions of ENU-treated PD-1^-/-^ mice. Granulomas are generally characterized as highly organized structures involving macrophages and T cells [26]. Here, CD3^+^ T cells and CD11b^+^ macrophages were confirmed to be present in the granulomas found in the liver of ENU-treated PD-1^-/-^ mice (Figure 2).

**Figure 2.**
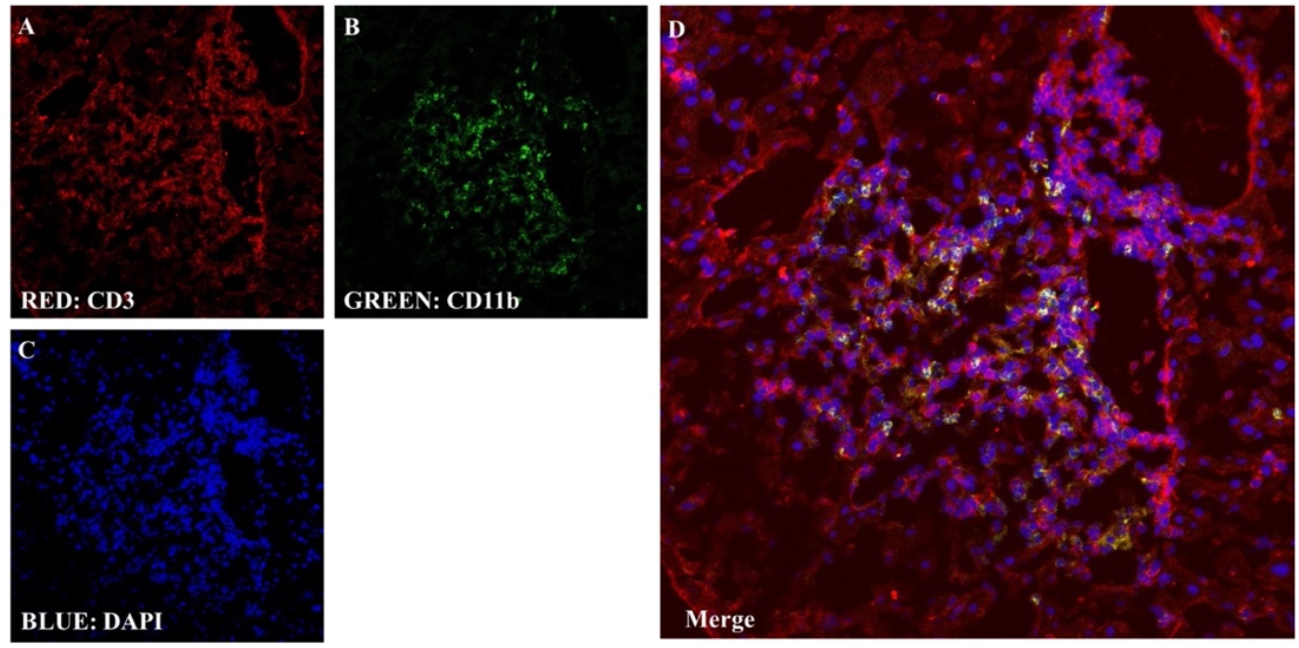
Cluster of lymphocytes and macrophages in granulomatous lesions found in the liver of ENU-treated PD-1^-/-^ mice. (**A-D**) Representative confocal microscopic images of a granuloma in the liver tissues from ENU-treated PD-1^-/-^ mice with an anti-CD3 antibody (**A**) and anti-CD11b antibody (**B**). Nuclei were counterstained with 4’,6-diamidino-2-phenylindole (DAPI) (**C**). Merge between CD3^+^ T cells (red), CD11b^+^ macrophages (green) and DAPI (blue) shows that granuloma comprises of both T cells and macrophages (**D**).

### The liver of ENU-treated PD-1^-/-^ mice begins to develop the granulomatous lesions at three to four months after birth

Different time points of ENU-treated mice were analyzed to study the stages of formation of the granulomatous lesions (Figure 3). Here, in the early stage, ENU-treated PD-1^-/-^ mice at three months after birth (one month after ENU treatment) did not show the formation of granulomatous lesions. Only lymphocytic foci were seen in the liver (Figure 3I-J). In contrast, the granulomatous lesions can be observed in the liver of PD-1^-/-^ mice even at five and seven months after birth (three and five months after ENU treatment) (Figure 3M-Q). This result shows that the manifestation of granuloma’s appearance does not occur instantly upon ENU treatment. Moreover, this result suggests that the mutation acceleration and accumulation occur in the genome of somatic cells, and the gradual chronic effect appears at a later stage of ENU treatment in the context of PD-1. In contrast, the ENU-treated PD-1^+/+^ mice also showed granuloma formation at seven months after birth (Figure 3G, Q). This result suggests the possible involvement of a chemical substance, ENU, that might be difficult to be eliminated and may trigger the formation of granulomatous lesions.

**Figure 3.**
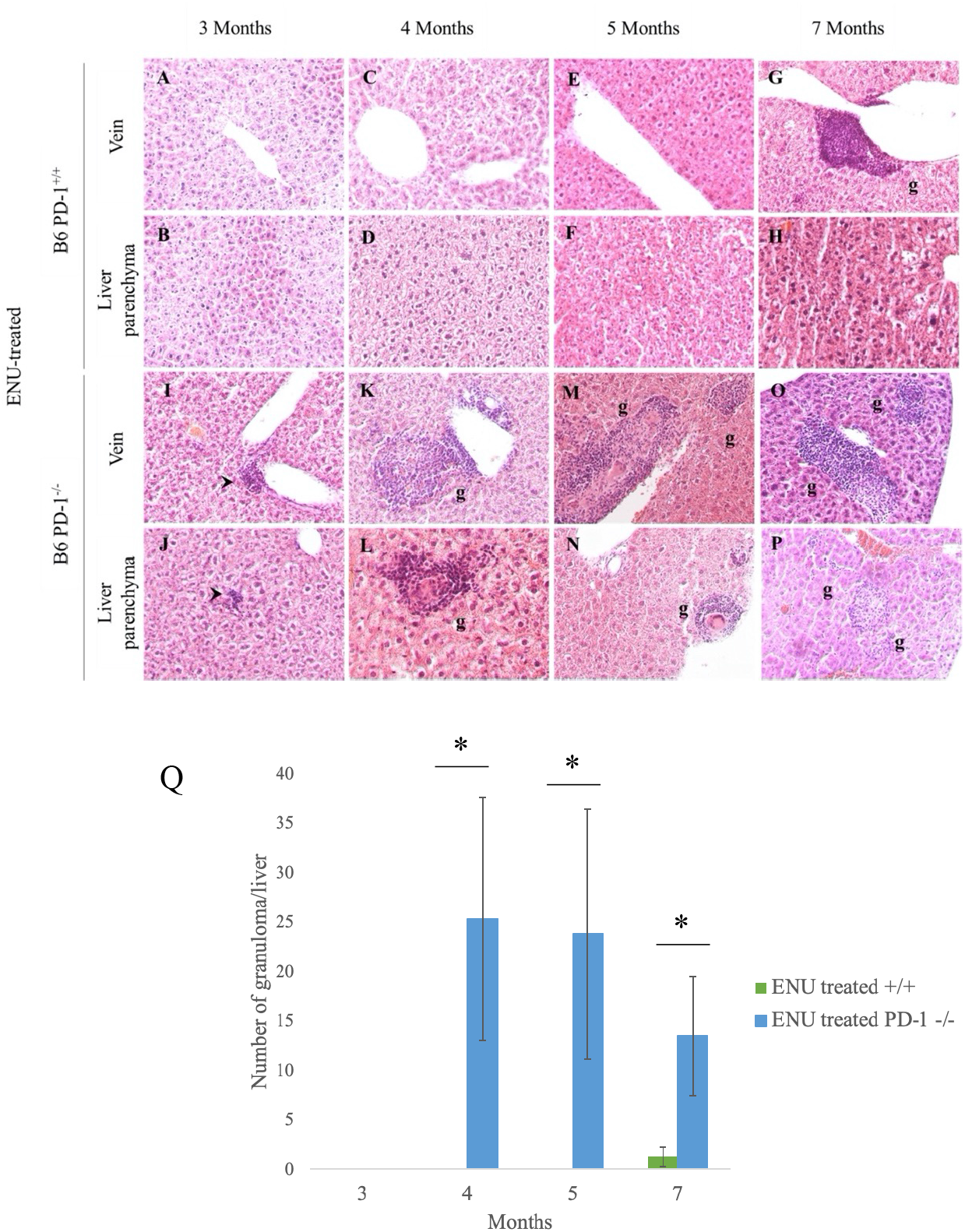
The liver of ENU-treated PD-1^-/-^ mice begins to develop the granulomatous lesions at three to four months after birth. (**A-P**) Representative histological sections of liver tissues from ENU-treated PD-1^+/+^ (**A-H**) and PD-1^-/-^ mice (**I-P**) at different time points after birth (one, two, three and five months after ENU treatment) are shown. Liver sections were examined by hematoxylin and eosin staining (200x magnification). At three months after birth, lymphocytic foci were seen in the liver tissue in both vein and parenchyma regions (one month after ENU treatment) (**I-J**). Granulomatous lesions were observed in the liver either around the vein or parenchyma regions from four months after birth (**K-P**) in ENU-treated PD-1^-/-^ mice and at seven months after birth (**G**) in ENU-treated PD-1^+/+^ mice. g(s) represent granulomas (**G, K-P**) and arrowheads indicate lymphocytic foci (**I-J**). Four (right lateral, left lateral, right medial, and left medial) liver lobes in every mouse were used for analysis (*N* ≥ 4 mice per ENU-treated group). (**Q**) Number of granuloma foci at different time points after birth (one, two, three, and five months after ENU treatment, *N* ≥ 4 mice per group). Data are expressed as means ± SEM. Statistical significance was determined by a two-tailed Mann-Whitney *U* test. *P<0.05.

### ENU-treated PD-1^-/-^ mice show the sign of hepatocytic necrosis

The array of histopathologic features in the ENU-treated mice was studied. Here, a few events of inflammatory lesions were observed upon examining the liver of ENU-treated mice. The sign of hepatocytic necrosis was observed in ENU-treated PD-1^-/-^ mice at three months after birth (Figure 4). It was also observed more frequently throughout the analyzed time points in ENU-treated PD-1^-/-^ mice (Figure 4E-I) than in ENU-treated PD-1^+/+^ mice (Figure 4B, C, I). The necrotic hepatocytes appear to be large and sometimes pale with swollen cytoplasm. In addition, adjacent lymphocyte infiltration was also found to surround these necrotic cells. The cells undergoing necrosis suggest the possibility of initiation of lymphocyte attack on the hepatocytes in the early stage of inflammation.

**Figure 4.**
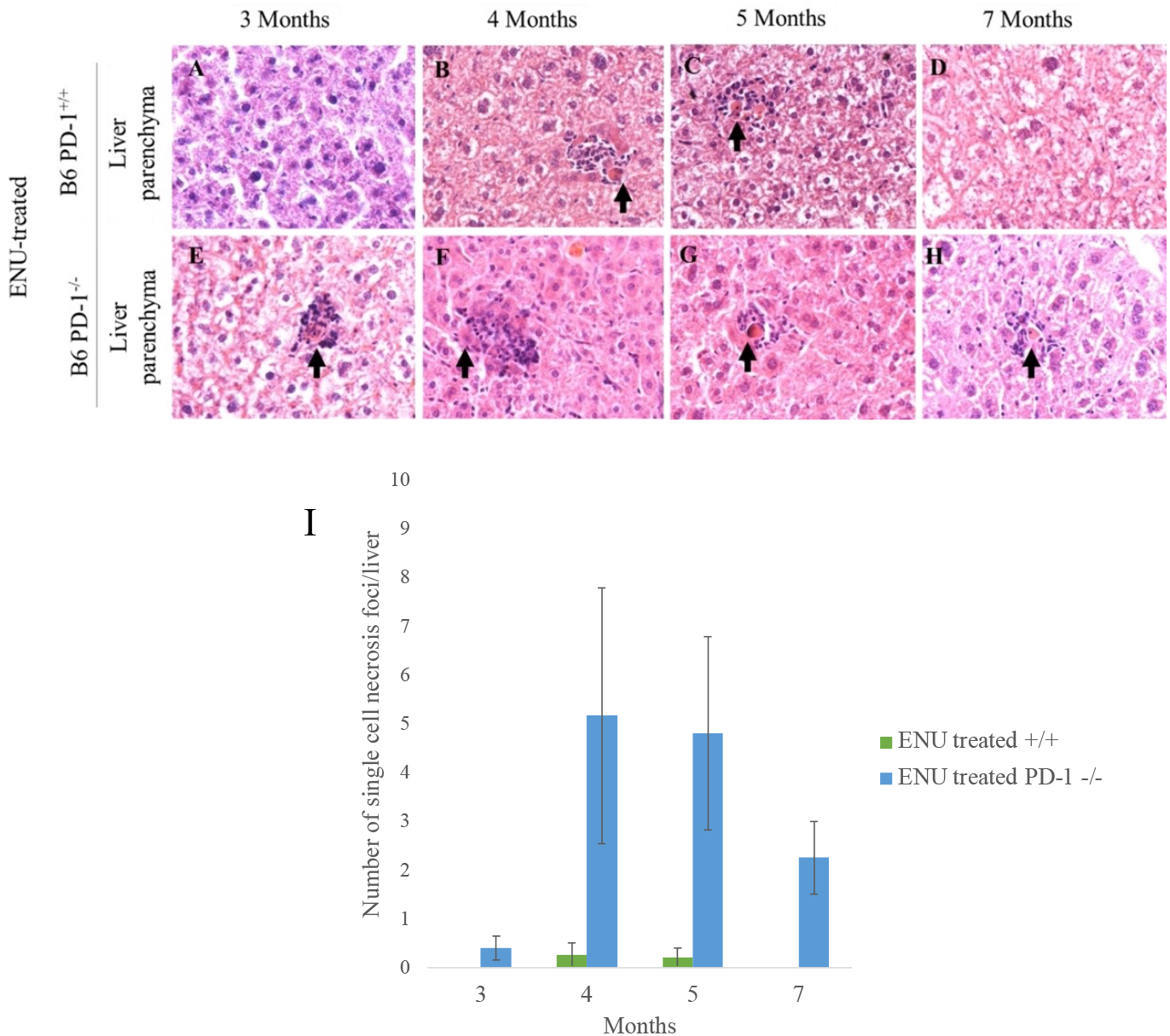
ENU-treated PD-1^-/-^ mice show the sign of hepatocytic necrosis. (**A-H**) Representative histological sections of liver tissues from ENU-treated B6 PD-1^**+/+**^ mice (**A-D**) and B6 PD-1^-/-^ mice (**E-H**) at different time points after birth (one, two, three, and five months after ENU treatment) are shown. Liver sections were examined by hematoxylin and eosin staining (400x magnification). The signs of dying hepatocytes surrounded by lymphocytes were observed in the liver parenchyma regions. Arrows indicate dying hepatocytes (**B,C,E,F,G,H**). Four (right lateral, left lateral, right medial, left medial) liver lobes in every mouse were used in the analysis (*N* ≥ 4 mice per ENU-treated group). (**I**) Number of single cell necrosis foci at different time points after birth (one, two, three, and five months after ENU treatment, *N* ≥ 4 mice per group). Data are expressed as means ± SEM.

### The liver of ENU-treated PD-1^-/-^ mice shows infiltration of mononuclear inflammatory cells

Another type of histopathologic feature that was most prevalently observed in liver tissues was the infiltration of mononuclear inflammatory cells. Lymphocytic foci were observed in all the tested ENU-treated PD-1^-/-^ mice (Figure 5). These mononuclear inflammatory cells appeared around the vein and in the liver parenchyma regions (Figure 5A-Q). In addition, mononuclear inflammatory cells were also found around the granulomatous lesions (Figures 1 and 3) and during the event of hepatocytic necrosis (Figure 4). These results suggest that lymphocytes are essential in regulating the granulomatous responses after the ENU treatment in the context of PD-1.

**Figure 5.**
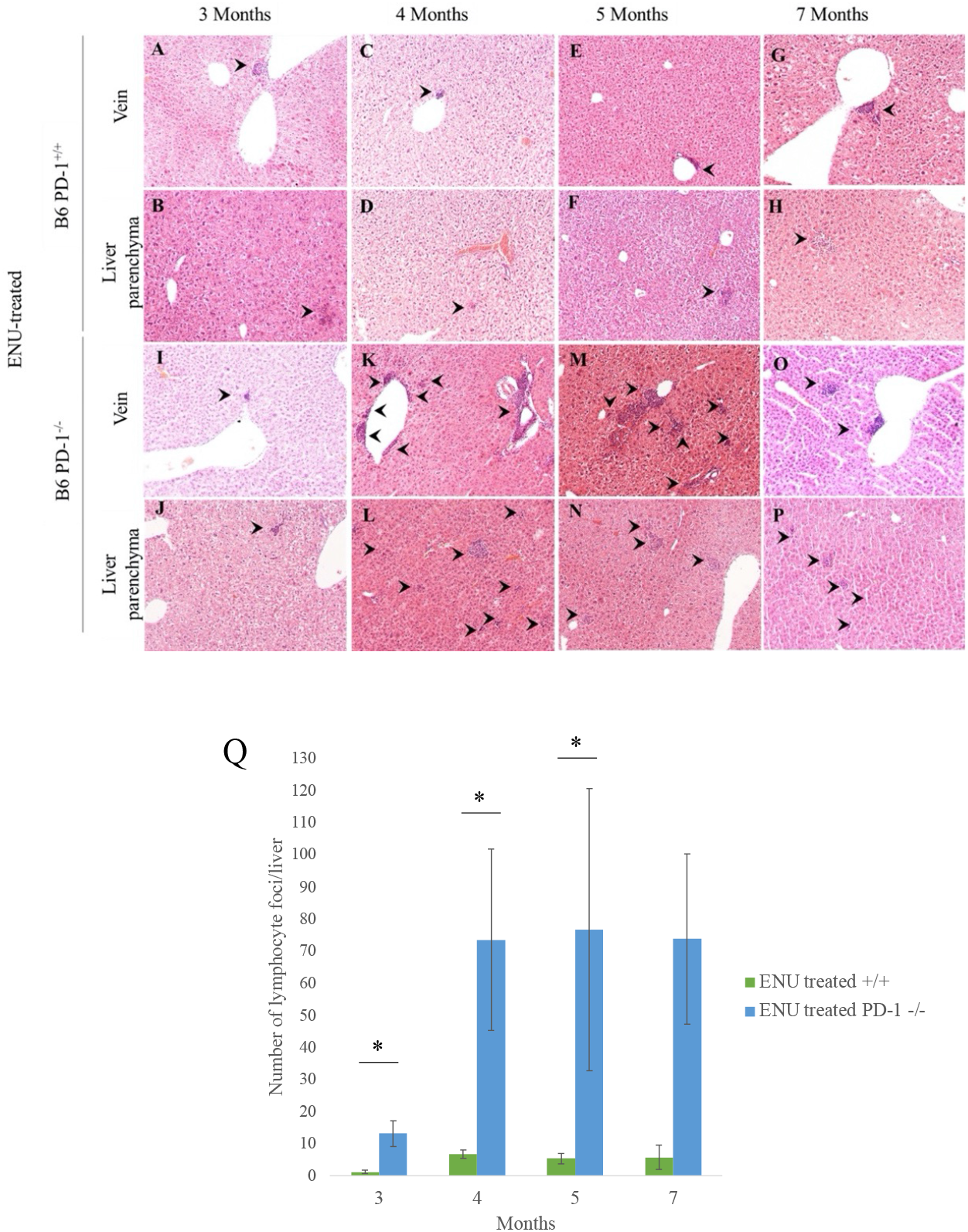
The liver of ENU-treated PD-1^-/-^ mice shows infiltration of mononuclear inflammatory cells. (**A-P**) Representative histological sections of liver tissues from ENU-treated PD-1^**+/+**^ mice (A-H) and PD-1^-/-^ mice (**I-P**) at different time points after birth (one, two, three, and five months after ENU treatment) are shown. Liver sections were examined with hematoxylin and eosin staining (100x magnification). Lymphocytic foci were frequently observed in both vein and parenchyma regions in the liver from two months after ENU treatment (**K-P**). Arrowheads indicate lymphocytic foci (**A-P**). Four (right lateral, left lateral, right medial, left medial) liver lobes in every mouse were used in the analysis (*N* ≥ 4 mice per ENU-treated group). (**Q**) Number of lymphocytic foci at different time points after birth (one, two, three, and five months after ENU treatment, *N* ≥ 4 mice per group). Data are expressed as means ± SEM. Statistical significance was determined by a two-tailed Mann-Whitney *U* test. *P<0.05.

### Quantitative assessment of inflammatory lesions in the liver of ENU-treated and untreated mice

We determine the total number of the inflammatory foci based on the signs of hepatocytic necrosis, lymphocyte infiltration, and granuloma formation. A total of four types of liver lobes were analyzed including right lateral, left lateral, right medial, and left medial lobes. The total number of inflammatory foci was higher in ENU-treated PD-1^-/-^ mice, compared to that of either ENU-treated PD-1^+/+^ mice, ENU-untreated PD-1^+/+^ mice, or ENU-untreated PD-1^-/-^ mice (Figure 6). These results strongly suggest that the ENU treatment accelerated the infiltration of inflammatory cells in the liver of mice, particularly when they lack the immuno-suppressive activity of PD-1.

**Figure 6.**
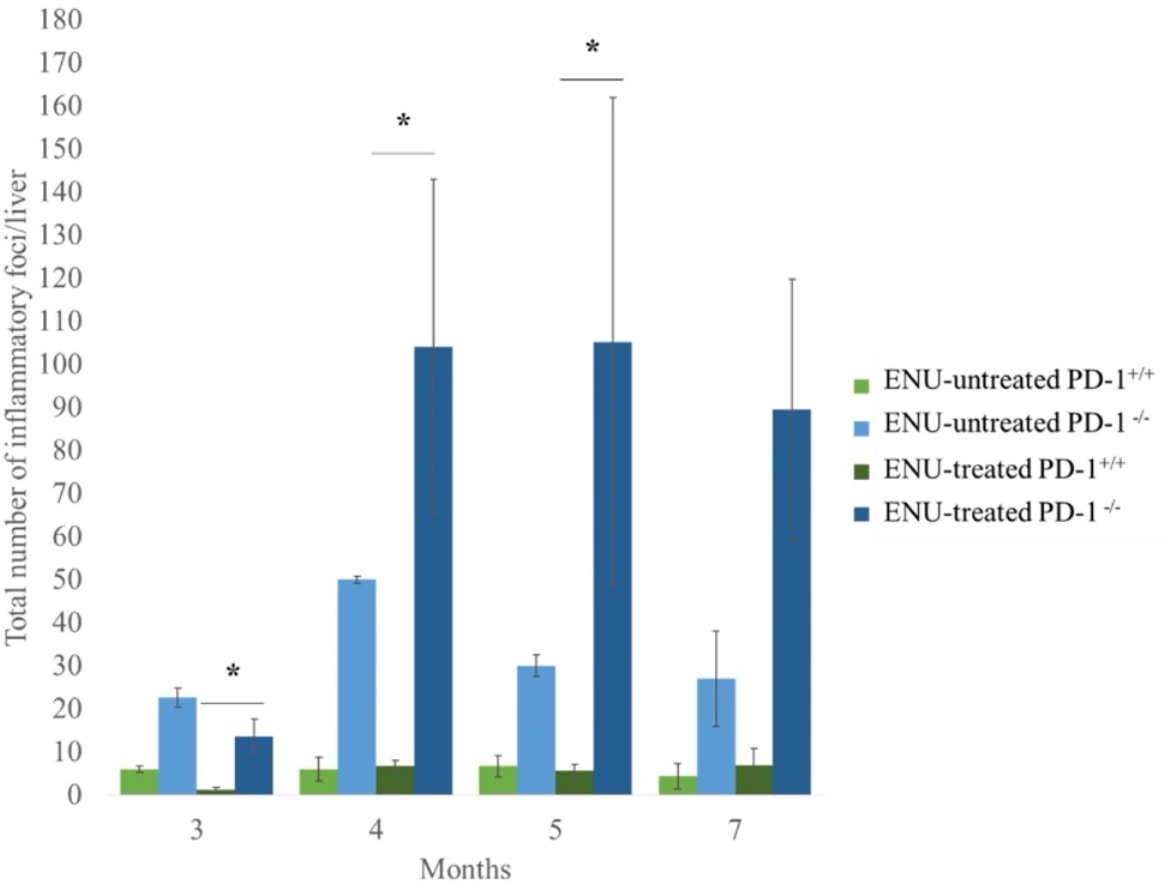
Quantitative assessment of inflammatory lesions in the liver of ENU-treated PD-1^+/+^ and PD-1^-/-^ mice. Number of inflammatory foci in ENU-untreated and ENU-treated mice (PD-1^+/+^ and PD-1^-/-^) at different time points after birth (one, two, three, and five months after ENU treatment, *N* ≥ 2 mice per ENU-untreated and *N* ≥ 4 mice per ENU-treated group). Data are expressed as means ± SEM. Statistical significance was determined by a two-tailed Mann-Whitney *U* test. *P<0.05.

### Histologic observations in organs other than the liver from three to five month after birth in ENU-treated PD-1^+/+^ and PD-1^-/-^ mice

Next, we investigated the inflammatory lesions in organs other than the liver in ENU-treated mice. Among the examined organs, the kidney, salivary gland, and pancreas showed lymphocytic focus formation in the ENU-treated PD-1^-/-^ mice (Figure 7). This could be due to the mutation accumulation in organs other than the liver, which has triggered the infiltration of mononuclear inflammatory cells.

**Figure 7.**
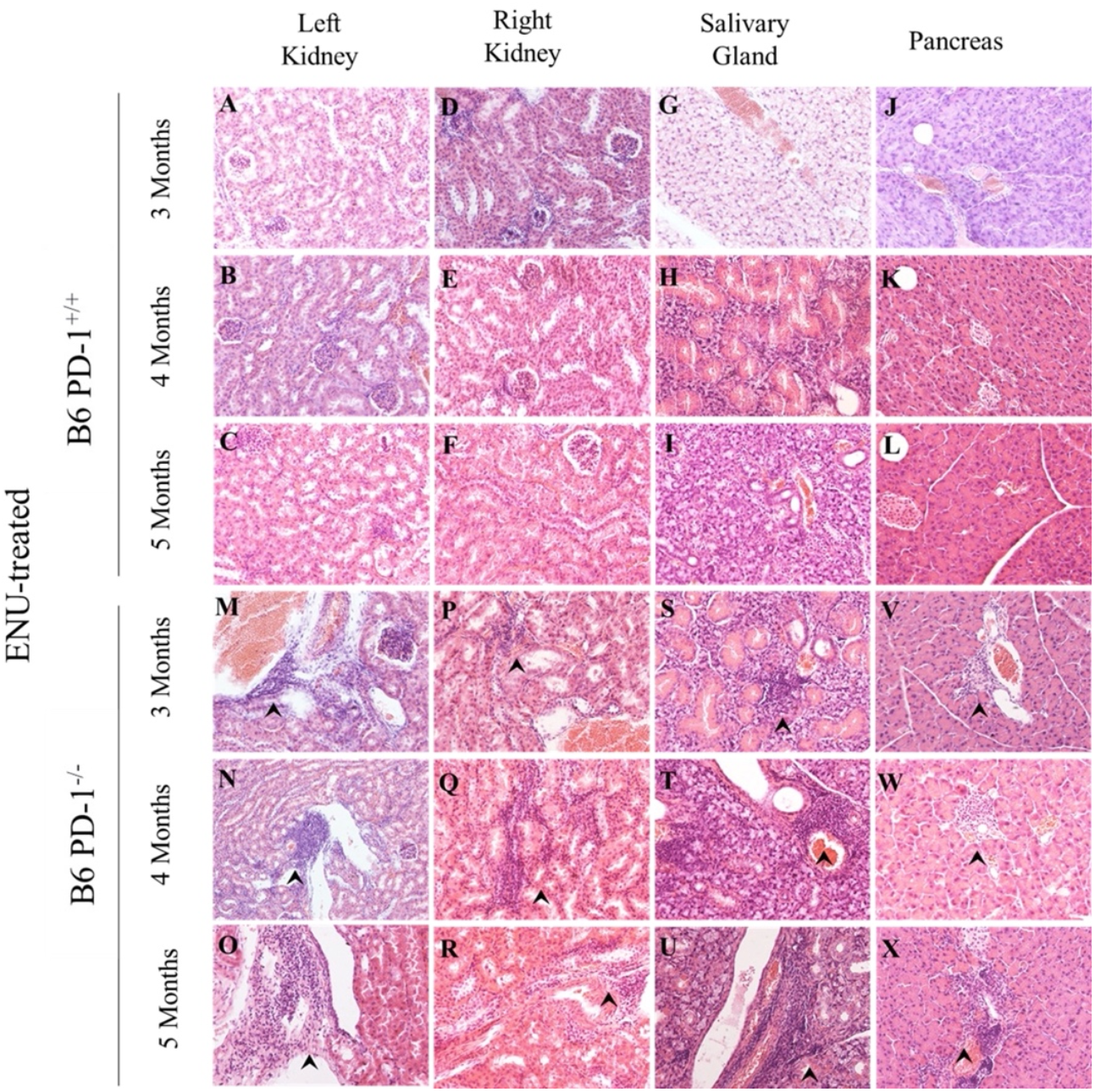
Histologic observations in organs other than the liver from three to five month after birth in ENU-treated PD-1^+/+^ and PD-1^-/-^ mice. (**A-X**) Representative histological sections of left kidney, right kidney, salivary gland, and pancreas from ENU-treated PD-1^+/+^ mice (**A-L**) and PD-1^-/-^mice (**M-X**) at different time points after birth (one, two, and three months after ENU treatment) are shown. All these tissue sections were examined by hematoxylin and eosin staining (100x magnification). Lymphocyte infiltration was observed in the left kidney (**M,N,O**), right kidney (**P,Q,R**), salivary gland (**S,T,U**), and pancreas (**V,W,X**) from three months after birth. Arrowheads indicate lymphocytic foci. Sections from left kidney, right kidney, pancreas, and salivary gland from every mouse were used for analysis (*N* ≥ 4 mice per ENU-treated group).

### The global absence of PD-1 and MSH2 in mice at seven months after birth induces the development of granulomas in the liver

To further examine the systemic effect of accumulation of genome mutations in somatic cells in the absence of PD-1, PD-1^-/-^MSH2^-/-^ mice were generated. The DNA mismatch repair (MMR) pathway plays a vital role in safeguarding the genome from DNA damage [15]. MMR acts on bases with mismatches, insertions, or deletions that are created during DNA replication and recombination, thereby inhibiting mutations in dividing cells. *MSH2* plays a pivotal role in repairing the base-base mispair by inserting and deleting nucleotide bases. Lack of *MSH2* promotes microsatellite instability (MSI), impaired DNA-damage response, and triple-repeat instability, and as a consequence leads to cancer development [15]. The MSH2^-/-^ mice have a shorter life span by exhibiting the development of lymphoma at an early age after birth [12,13,14]. Since tumors lacking the MMR function can induce good clinical responses upon PD-1 blockade in human patients [16], it would be intriguing to see the phenotype(s) of mice, in which normal somatic cells without the MMR activity are kept under surveillance by immune cells lacking the PD-1 checkpoint molecules.

Interestingly, at seven months after birth, granulomatous lesions were also observed in the liver of PD-1^-/-^MSH2^-/-^ mice (Figure 8). Induction of mutation accumulation by the deletion of *MSH2* appeared to result in the development of progressive chronic inflammation through the manifestation of granulomatous lesions especially in the absence of immuno-suppressive activity of PD-1 in mice.

**Figure 8.**
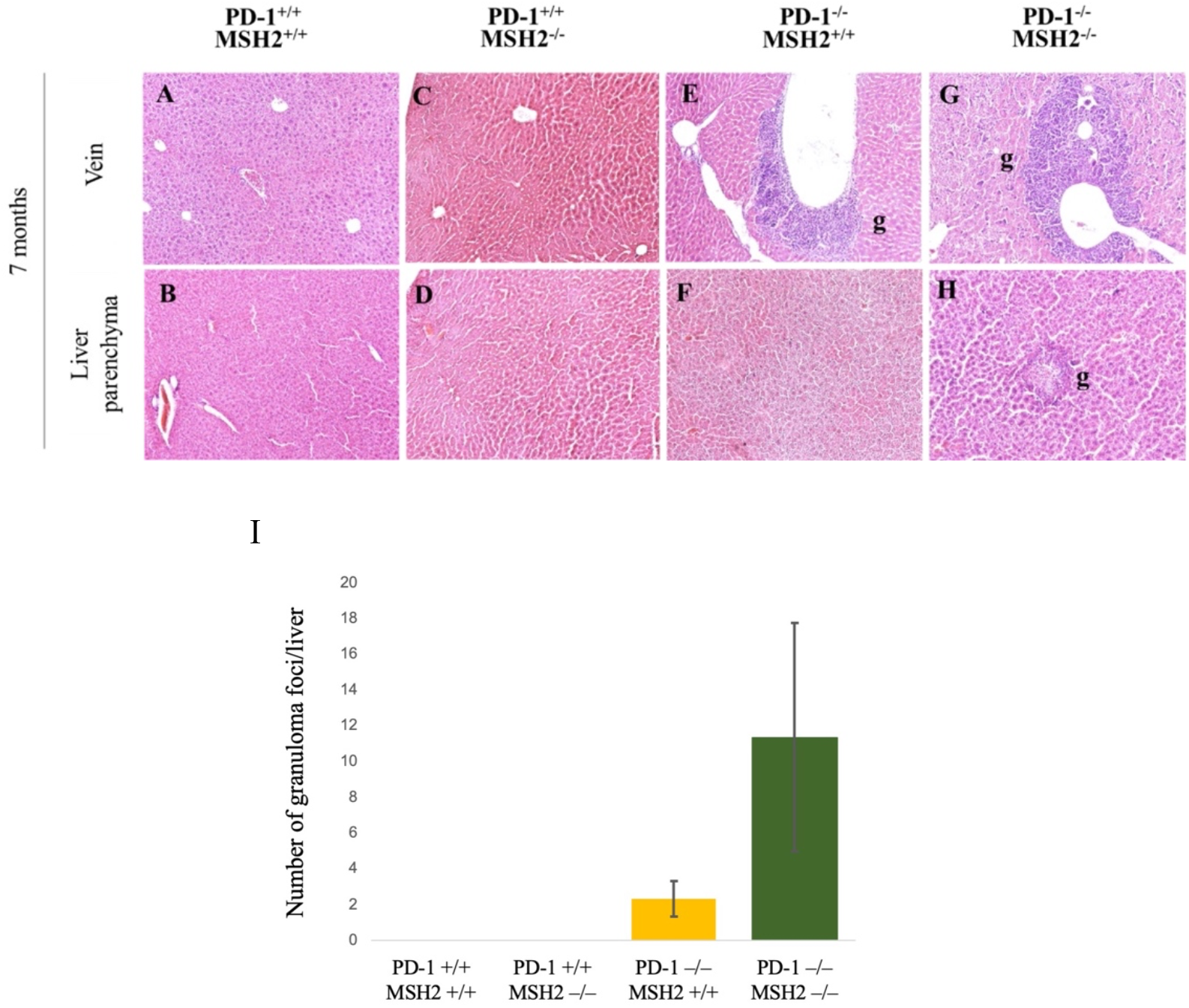
The global absence of PD-1 and MSH2 in mice at seven months after birth induces the development of granulomas in the liver. (**A-H**) Representative histological sections of liver tissues from PD-1^+/+^MSH2^+/+^, PD-1^+/+^MSH2^-/-^, PD-1^-/-^MSH2^+/+^, and PD-1^-/-^ MSH2^-/-^ mice at seven months of age are shown. Liver sections were examined by hematoxylin and eosin staining (100x magnification). Granulomas were seen in both vein (**G**) and parenchyma (**H**) regions in the liver of PD-1^-/-^MSH2^-/-^ mice. g(s) represent granulomas (**E,G,H**). (**I**) The bar graph showed the number of granulomas in PD-1^+/+^MSH2^+/+^, PD-1^+/+^MSH2^-/-^, PD-1^-/-^MSH2^+/+^, and PD-1^-/-^MSH2^-/-^ mice. Four (right lateral, left lateral, right medial, and left medial) liver lobes in every mouse were used in the analysis. Numbers of mice analyzed were: 4 (PD-1^+/+^MSH2^+/+^), 2 (PD-1^+/+^MSH2^-/-^), 6 (PD-1^-/-^MSH2^+/+^), and 6 (PD-1^-/-^MSH2^-/-^). Data are expressed as means ± SEM.

## Discussion

Symptoms of mild autoimmunity developed gradually only after PD-1^-/-^ mice (on the B6 genetic background) got old [4]. In a model experiment of cancer immunotherapy, PD-1 was shown to prevent cytotoxic T cells that could recognize and respond to mutant antigens (in the context of a class I MHC molecule) from attacking cancer cells with the cognate genome mutations [6]. The larger number of genome mutations in cancer cells led to the more robust immune responses against such cells after the PD-1 blockade [7]. In order to understand the common molecular basis for these findings, we hypothesize that we might have acquired PD-1 during evolution in order to avoid/suppress autoimmune reactions against neoantigens originated from gradually accumulating random genome mutations in our aged somatic cells [8].

Young PD-1^-/-^ mice on the B6 genetic background usually do not show any particular signs of autoimmunity [4]. This is probably because they still have not accumulated a significant number of genome mutations in their somatic cells. Aged B6 PD-1^-/-^ mice, on the other hand, might have accumulated a relatively large number of mutations in the genome of their somatic cells, and in the absence of PD-1, the immune system could misrecognize them as ‘nonself’ components that are to be cleared from the mouse bodies.

Technically, it is still challenging to assess the mutation rate (the number of mutations per cell per year) in each of normal (untransformed) somatic cells because of their enormous diversity and heterogeneity [17]. It becomes feasible only when DNA is extracted from an almost clonal cell population with the same ancestral origin, like the cells from a single crypt in the intestine, and analyzed. According to Cagan et al. [18], the nucleotide-substitution rates per genome per year were 47 and 796 in humans and mice, respectively (∼17 fold difference between the species), while the indel rates per genome per year were 2.5 and 158 in humans and mice, respectively (∼63 fold difference between the species).

To test the above hypothesis about the physiological function of PD-1, we tried in this study to induce the accumulation of mutations in the genome of somatic cells in young PD-1^-/-^ and PD-1^+/+^ mice by employing two different methods of random genome mutagenesis, (i) administration of a potent mutagen ENU into the peritoneum of mice and (ii) deletion from the mouse whole body of *MSH2*, an essential mismatch-repair gene. Interestingly, formation of the granulomatous lesions was observed in both of the mutation-promoting mouse models, especially in the absence of the immuno-suppressive activity of PD-1. These results suggest that some neoantigens originating from the mutated genome could be presented on self-MHC molecules to induce T-cell recognition and activation, but PD-1 usually suppresses immune responses of T cells against such molecular determinants. In the absence of PD-1, however, T cells recognizing the neoantigen/self-MHC complexes are unleashed, and pathological immune reactions are elicited [8].

Recently, Damo et al. [19] expressed the lymphocytic choriomeningitis virus (LCMV)-derived antigenic epitopes only in the skin of their transgenic mice in a strictly regulated/inducible way, by using an elegant genetic strategy. They showed that PD-1 is responsible for the maintenance of the CD8 T-cell tolerance towards peripherally/restrictedly expressed neoantigens, also supporting the above hypothesis of ours about the physiological function(s) of PD-1. It would be important to note that our random mutagenesis strategy shown here creates only the peripheral (*i.e*., somatic, not germline) genome mutations that cannot be found in the thymus. In this sense, we have also been focusing on the role(s) of PD-1 in the induction of peripheral immunological tolerance by employing simple random-mutagenesis experiments.

In the case of ENU-treated PD-1^-/-^ mice in our study, the main inflammatory lesions were observed in the liver. This could be due to the site of ENU injection in this study, *i.e*., inside the peritoneal cavity. Chemical substances injected intraperitoneally should be absorbed into many capillaries in the peritoneum and carried into mesenteric veins [20]. Mesenteric veins then converge into the portal vein, delivering the abdominally absorbed substances to the liver [21]. Therefore, it has been speculated that the accumulation of the ENU molecules may have been highly concentrated in the liver cells, almost selectively and preferentially altering the genome of hepatic cells to create the ‘neo-self’ components (neoantigens) there to initiate immune responses.

On the other hand, the main inflammatory lesions were also observed in the liver of the PD-1^-/-^MSH2^-/-^ mice. Although it is conceivable that the genome of liver cells could be one the main targets of mutagenesis caused by the *MSH2* deletion, tumor development in the liver of the MSH2^-/-^ mice is a relatively rare event [12,13]. Therefore, we assume it is more likely that the systemic and comparable level of genome mutagenesis actually takes place throughout the body of the MSH2^-/-^ mice, but there could be some unknown reasons for the preferential development of the inflammatory lesions in the liver of the knockout mice, especially under the decreased activity of PD-1. The same logic could also be applicable for the inflammatory changes mainly observed in the liver of the ENU-treated PD-1^-/-^ mice. Interestingly, in the human clinical settings, the liver is one of the most frequently targeted organs of the immune-related adverse events (IRAEs) in cancer patients administered with the immune-checkpoint inhibitors (ICIs) like nivolumab or pembrolizumab [22,23, 24,25]. An interesting feature of inflammatory changes that are commonly observed in the ENU-treated PD-1^-/-^ mice and the PD-1^-/-^MSH2^-/-^ mice is the development of granulomas in the liver. Granuloma formation occurs when the host fails to eliminate antigens from infectious microbes, or when inflammatory disorders last over a long period of time [26]. Granulomatous inflammation can thus be classified as a form of chronic inflammation characterized histologically by an aggregate of activated macrophages (often called epithelioid cells) surrounded by lymphocytes [26]. The liver is a unique organ with a large number of Kupffer cells (long-lived tissue macrophages) scattered and attached to the luminal face of the sinusoids [27]. Recently, it was shown that the activation status of macrophages is also negatively regulated by PD-1 [28].

Therefore, it would be possible to assume that the threshold for the activation of Kupffer cells is lowered by the absence of PD-1, and together with the hyperactivated nature of T cells without PD-1, subtle changes in the amino-acid sequences of proteins, originated from random genome mutations in the liver, induced the neoantigen presentation by Kupffer cells to T cells and subsequently elicited the development of granulomatous inflammations in the organ.

It is also a fascinating idea to assume that a combination of the increased number of random genome mutations in somatic cells and the decreased activity of PD-1 in T cells and tissue-resident macrophages may somehow result in the development of at least a part of granulomatous inflammatory diseases with an unknown etiology, in which sarcoidosis [29], Crohn disease [30], and granulomatosis with polyangiitis (GPA) [31] would be included.

In the settings of cancer immunotherapy, some mutations in the protein-coding regions in the genome of cancer cells might result in the production of neoantigens, and a portion of them could be successfully presented in the context of MHC molecules on the cell surface and recognized by T cells as the ‘nonself’ components [6]. However, when PD-1 suppresses harmful immune responses against normal somatic cells harboring neoantigens encoded by genome mutations in aged individuals, the beneficial immunity against neoantigens generated in cancer cells also should be suppressed by PD-1 [8].

Upon PD-1 blockade, an immune reaction against neoantigens originating from genome mutations would be generally unleashed. This should probably be one of the central molecular mechanisms for the IRAEs found in cancer patients with checkpoint-blockade immunotherapy [32,33]. Importantly, however, the number of somatic mutations in the genome of cancer cells is usually much larger than that in normal somatic cells. Therefore, from the immunological point of view, it would be natural for us to observe the stronger immune reactions against cancer cells than toward normal somatic cells upon cancer immunotherapy with PD-1 blockade [8].

## Conclusion

We hypothesize that we might have acquired PD-1 during evolution in order to avoid/suppress autoimmune reactions against neoantigens derived from natural and inert mutations in the genome of aged individuals. To test the hypothesis, we introduced random mutations into the genome of somatic cells in young PD-1 KO mice. We employed two different procedures of random mutagenesis: administration of a potent chemical mutagen ENU into the peritoneal cavity of mice and deletion from the mouse whole body of *MSH2*, which is essential for the suppression of accumulation of random mutations in the genome. We observed granulomatous inflammatory changes in the liver of the ENU-treated PD-1 KO mice, but not in the wild-type counterparts. Such lesions also developed in the PD-1/MSH2 double KO mice, but not in the MSH2 single KO mice. These results support our hypothesis shown above.

## Acknowledgments

We thank Tasuku Honjo (Kyoto University) and Teruhisa Tsuzuki (Kyushu University) for providing us with the PD-1 KO mice and MSH2 KO mice, respectively. We also thank lab members in NAIST and Mizuki Ohno (Kyushu University) for valuable discussion. This study was supported by JSPS KAKENHI Grant-in-Aid for Scientific Research B (21H02717) and for Challenging Research (Exploratory) (22K19508); ONO Pharmaceutical Co., LTD.

## Author contributions

I. Nagaretnam performed the mouse experiments, analyzed the data, and wrote the manuscript. S. Suzuki, A. Yoneshige, F. Takeuchi and A. Ito supported the histological and pathological analyses. A. Ozaki and M. Tamura performed the ENU-administration experiments. T. Shigeoka analyzed the data. Y. Ishida conceived the project, acquired the research grants, supervised the research, and wrote the manuscript.

